# Gut-resident C. perfringens impedes rotavirus vaccine efficacy

**DOI:** 10.1101/2024.06.17.599343

**Authors:** Vu L. Ngo, Yanling Wang, Zhenda Shi, Sasirekha Ramani, Baoming Jiang, Andrew T. Gewirtz

## Abstract

**Background & Aims:** The extent to which live orally-administered rotavirus (RV) vaccines elicit protective immunity is highly heterogeneous. We hypothesized microbiota composition might influence vaccine efficacy.

**Methods:** We tested this concept by examining extent to which colonizing mice with segmented filamentous bacteria (SFB) influenced RV vaccine efficacy.Influence of human microbiomes on RV vaccination was studied via administering germ-free mice fecal microbial transplants (FMT) from children with robust or minimal RV vaccine responsiveness. Post-FMT, mice were subjected to vaccination and challenge doses of RV.

**Results:** SFB administration resulted in a phenotype reminiscent of RV vaccine failure, i.e. minimal generation of RV antigens and, consequently, lack of anti-RV antibodies resulting in proneness to RV challenge once SFB levels diminished. Transplant of microbiomes from children to mice recapitulated donor vaccination phenotype. Specifically, mice receiving FMT from high-responding children exhibited high levels of fecal RV antigen shedding and RV antibodies in response to RV vaccination and, concomitantly, were impervious to RV challenge. In contrast, mice receiving FMT from children who had not responded to RV vaccination exhibited only modest responses to RV challenge and, accordingly, remained prone to RV challenge. Microbiome analysis ruled out a role for SFB but suggested that RV vaccine failure might involve *Clostridium perfringens*. Oral administration of cultured *C. perfringens* to gnotobiotic mice partially recapitulated the RV vaccine non-responder phenotype. Analysis of previously-reported microbiome data found C. perfringens abundance in children associated with RV vaccine failure.

**Conclusion:** Microbiota composition influences RV vaccine virus infection and, consequently, protective immunity. *C. perfringens* may be one, perhaps of many, bacterial species harbored in the intestine of RV-vaccine non-responders that influences RV vaccine outcomes.

## INTRODUCTION

RV vaccines have markedly reduced the RV disease burden in high-income countries but have proven less effective some low-income countries as assessed by both their impact upon disease burden and the extent to which they elicit anti-RV antibodies in vaccinees ^1, 2^. RV vaccines are live attenuated viruses and thus, like virulent RV, must infect their host in order to elicit protective immunity. Failure of RV vaccines to elicit anti-RV antibodies in individual vaccinees associates with the absence of a transient post-vaccination increase in fecal RV antigens thus strongly suggesting that some instances of RV vaccine failure reflect inability of RV vaccine viruses to infect their hosts ^3^. On the hand, one could envisage that such instances might not be problematic in that individuals harboring microbiota that prevent RV infection would not be in need of an RV vaccine. However, one can imagine scenarios in which the protection provided by gut microbiota prevents infection of the vaccine virus but is not robust enough to prevent infection by more virulent circulating RV strains. Furthermore, if the presence of the microbiota that impeded infection of the RV vaccine was transient, such individuals would lack protection against virulent RV and thus potentially become examples of RV vaccine failure.

An example of a transient microbiota resident impeding RV infection is that of segmented filamentous bacteria (SFB) ^4^. Specifically, mice naturally colonized, with or exogenously administered, SFB are resistant to RV infection, particularly at early time points following SFB exposure when its levels are highest. Herein, we examined the extent to which SFB and/or microbes present in humans exhibiting poor antibody responses to RV vaccination impacted efficacy of RV vaccination in mice. We indeed observed that administration of SFB to mice resulted in an RV vaccine failure phenotype supporting our general hypothesis. Investigation of the extent to which this finding might apply to humans led us to study fecal microbes in a group of Mexican children RV vaccinees. SFB was not detected thus precluding it influencing RV vaccine responsiveness in this cohort. However, conceptual relevance was evident in that transplant of fecal microbes from RV vaccine non-responders to germfree mice recapitulated the RV vaccine failure phenotype. This phenotype could be partially mimicked by colonization of mice with Clostridia Perfringrens, which was present in one of the RV vaccine non-responders. These results support the concept that microbiota composition can influence RV vaccine responsiveness and that C. pefringrens is one example of a human microbiota constituent capable of promoting RV vaccine failure.

## RESULTS

### Administering mice SFB results in an RV vaccine failure phenotype

Infection of adult mice with RV has similarities to RV vaccination in humans in that, in both cases, a live virus asymptomatically infects the intestine leads to the transient presence of RV antigens that results in generation of anti-RV antibodies that help clear the virus and provide lasting protection against future RV infection ^5, 6^. One microbe capable of impeding RV infection in mice is segmented filamentous bacteria (SFB), which is frequently present in some but by no means all mouse colonies ^4^. SFB levels can be persistently high in some colonies of mice with genetically-encoded immune deficiencies resulting in stark lasting resistance to RV infection. In contrast, exposure of WT mice to SFB in young adulthood, i.e. 4-8 weeks of age, results in transiently high SFB levels and, consequently, temporary resistance to RV infection.

Accordingly, we envisaged that having relatively high levels of SFB, or SFB-like bacteria at the time of RV vaccination might be a possible cause of RV vaccine failure. To interrogate this hypothesis, mice were orally administered a fecal suspension that lacked or contained SFB one week prior to a low-dose orally administered RV inoculation. Mice administered FMT lacking SFB exhibited a week-long course of fecal RV antigen shedding and displayed anti-RV antibodies in serum and feces on day (d) 21 and 28 post-RV inoculation **(Figure 1)**.

**Figure 1.**
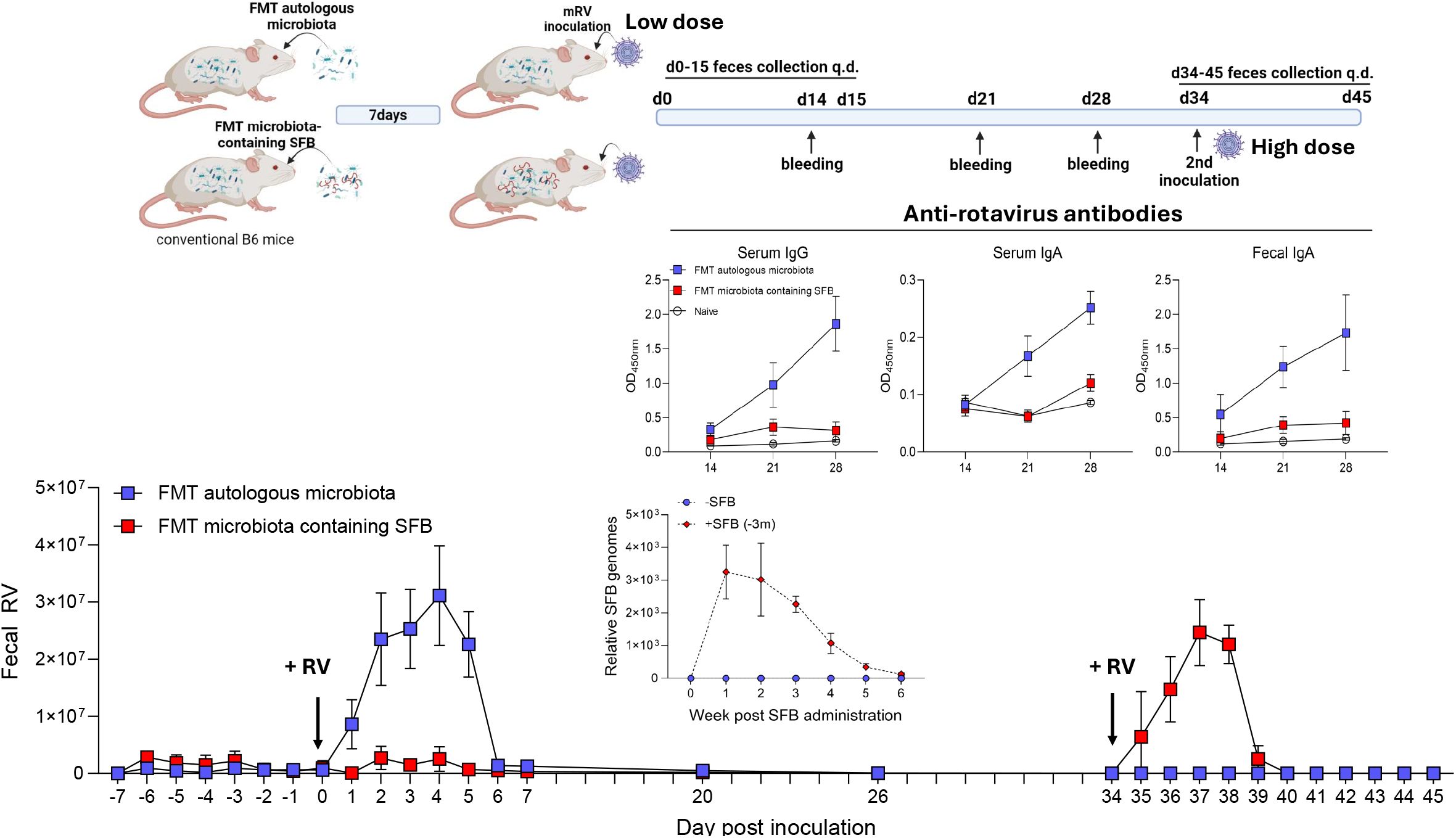

Concomitantly, such mice were protected against re-infection as evidenced by the lack of fecal antigens following a d34 high-dose RV challenge. In contrast, mice receiving an SFB-positive FMT displayed few fecal RV antigens and generated only very modest levels of anti-RV antibodies, which were not sufficient to protect against RV infection as evidenced by RV fecal antigen levels following the d34 high-dose RV challenge. Such susceptibility to the d34 RV infection may have reflected the higher viral dose and/or that SFB levels had peaked within 2 weeks exposure and were very low by day 34 and thus not sufficient to protect against RV challenge. Regardless, these results serve as proof of principle that select microbiota can impede infection of RV vaccine to result in lack of protection against future RV challenge.

### Transplant of children’s fecal microbiomes to germfree recapitulates RV vaccine responsiveness

The extent to which SFB is present in humans is not entirely clear although it has been detected in some come cohorts/individuals ^7, 8^. In any case, we view our work on mouse studies of SFB as an exploration of concept rather than relating to a specific role for SFB in RV vaccine failure per se. Thus, we sought to extend our studies to humans not by probing for SFB per se but by broadly investigating the extent to which microbiotas of children exhibiting poor antibody responses to RV vaccination might harbor bacteria capable of impeding RV infection and anti-RV antibody generation. We leveraged previously-collected fecal samples linked to previously determined measures of post-vaccination anti-RV antibody titers from a previous study of Mexican infants administered the RotaRix vaccine ^9^. Fecal samples from the infants exhibiting the highest and lowest post-vaccination anti-RV titers were selected for transplant into groups of germfree C57BL/6 mice **(Figure 2)**. Mice were then administered vaccination and challenge doses of RV 7d and 34d post-FMT, respectively. Fecal RV antigens and anti-RV antibodies were monitored as above. Mice receiving FMT from the non-responding infant exhibited a phenotype reminiscent of RV vaccine failure. Specifically, relative to the mice receiving FMT from the high-responding infant, the exhibited low post-vaccine levels of fecal RV antigens and anti-RV antibody titers rendering them somewhat susceptible to RV infection from the d34 inoculation. We repeated this experiment using fecal samples from infants with the second highest and lowest post-vaccination titers and observed a similar pattern of results. No recipients of either FMT displayed detectable levels of SFB. These results support the notion that microbes harbored by select individuals contribute to infection and, consequently, efficacy of RV vaccines.

**Figure 2.**
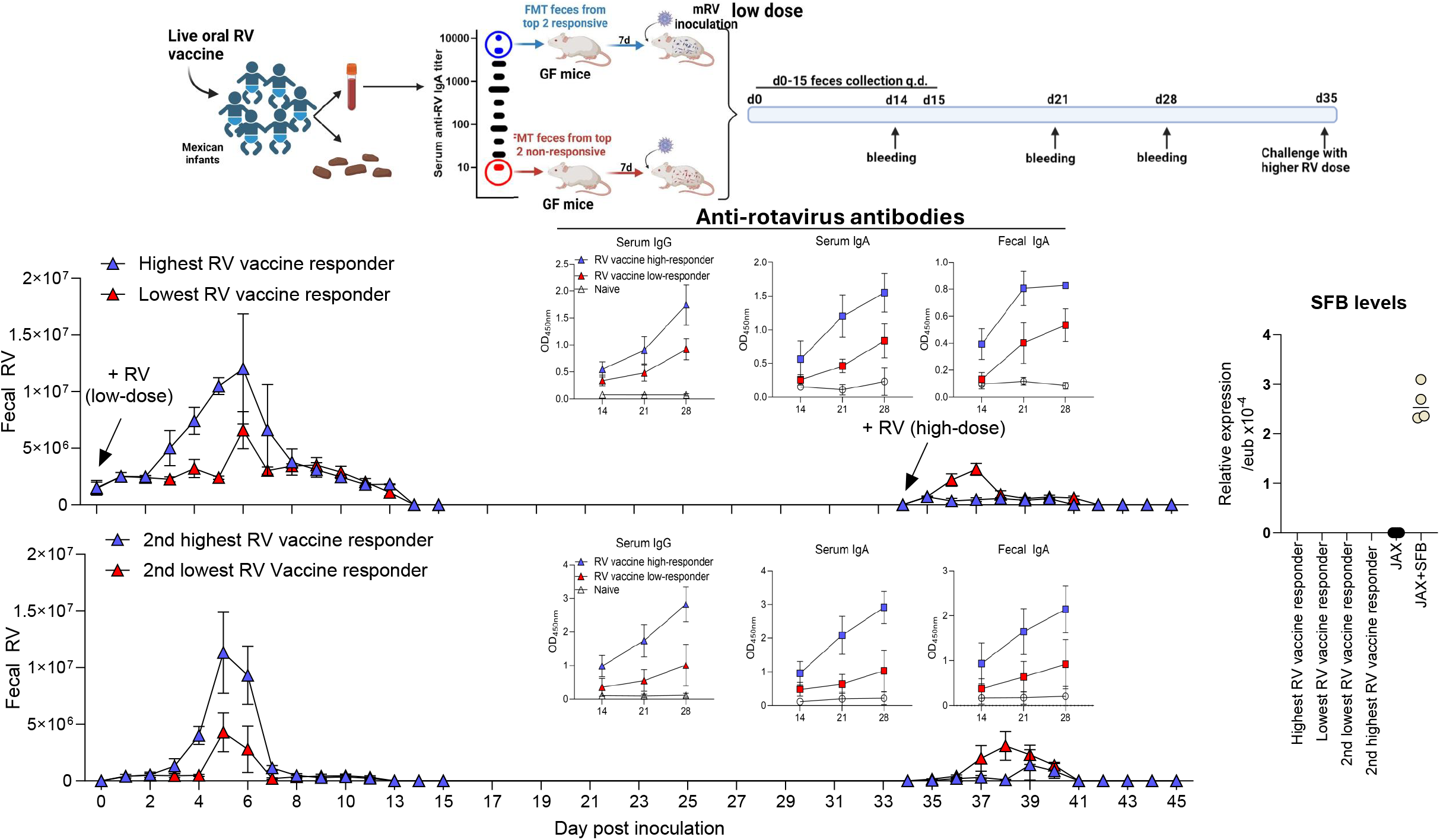

### C. perfringens, present in an RV vaccine non-responder impedes antibody response to RV vaccination

We next sought to identify bacteria that had engrafted in the FMT recipients potentially enabling them to influence outcomes of RV exposure. Hence, we analyzed fecal microbiomes of FMT recipients by 16S sequencing 7d post-transplant, i.e. at the time of RV inoculation **(Figure 3)**. Principle coordinate analysis showed a clear difference in overall composition was evident between mice receiving the FMT from the top responder and non-responder although overall a-diversity (richness) was surprisingly low in both groups of recipients, potentially reflecting the fecal samples were severely dehydrated and thus may have contained relatively few viable microbes (we lacked sufficient sample to direct analyze donor samples but, in any case, that would not have indicated microbe viability). This caveat notwithstanding, we hypothesized that taxa present in the non-responder recipients may nonetheless be specific examples of bacteria that are present in some RV vaccine non-responders and, furthermore, capable or promoting RV vaccine failure. Accordingly, we considered the possibility that Clostridia Perfringrens, which comprised 10-20% of the 16S reads in the responder FMT recipients, and was not detected in the non-responder recipients, might be capable of promoting RV vaccine failure.

**Figure 3.**
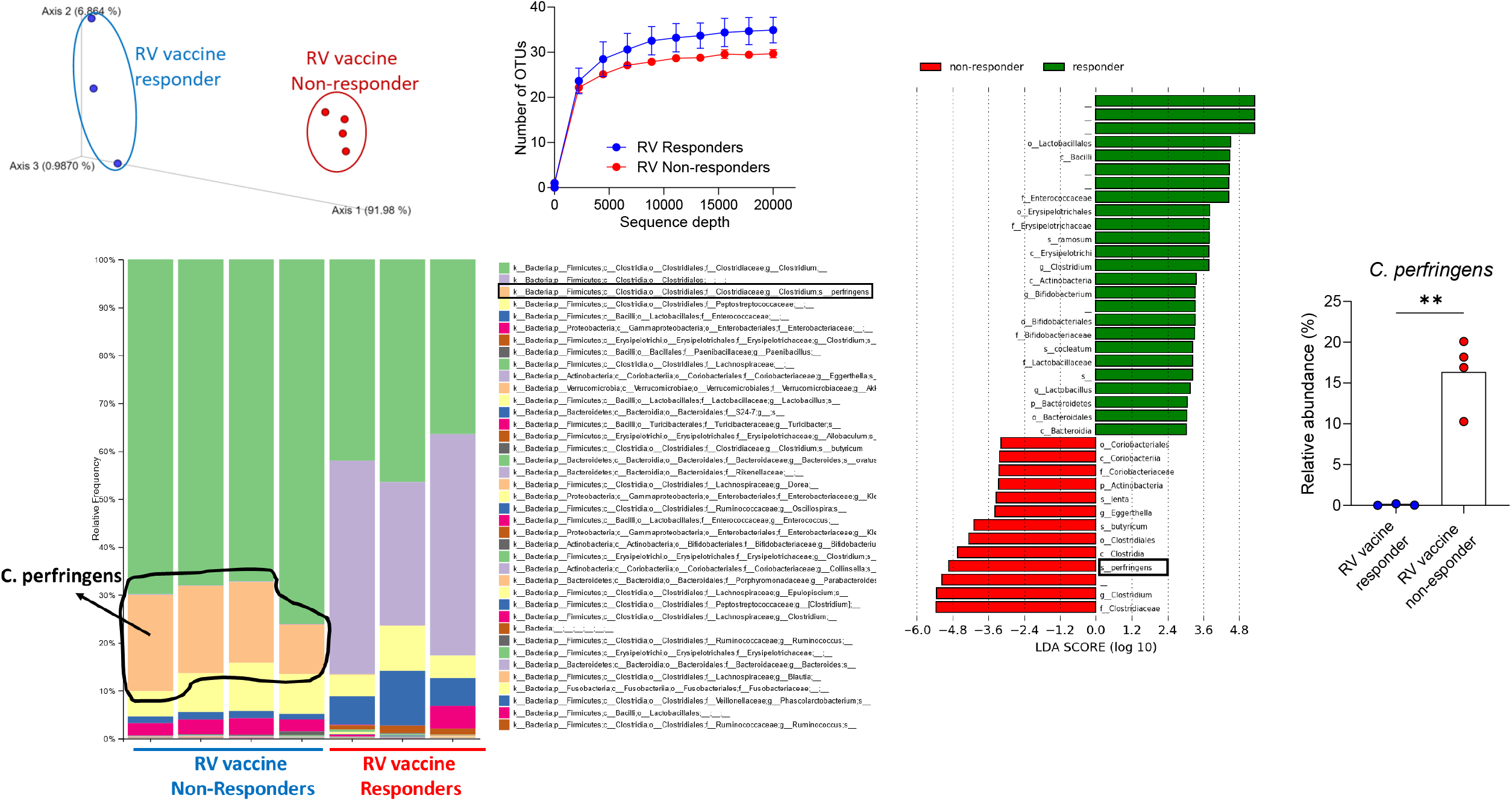

We first administered a human clinical C. pefringrens isolate to conventional mice one week prior to RV inoculation. Such C. pefringrens was not detectable in feces at the time of RV administration and resulted in a modest delay in peak levels of fecal RV shedding but did not impact generation of serum or fecal anti-RV antibodies **(Figure 4)**. Considering this results in the context of our FMT results, in which C. pefringrens seemed to readily colonize the gut, we reasoned it may reflect inability of C. perfringens to outcompete resident mouse microbiota. We sought to overcome this hurdle via use of mice with a minimal microbiota, namely mice colonized the 8-species referred to as the Altered Schaedler Flora (ASF) ^10^. ASF mice display minimal colonization resistance but yet lack many of the overt mucosal immunologic abnormalities of germfree mice. Indeed, C. pefringrens was readily detected in feces of ASF mice 7d following its administration, at which time mice were inoculated with RV **(Figure 5)**.

**Figure 4.**
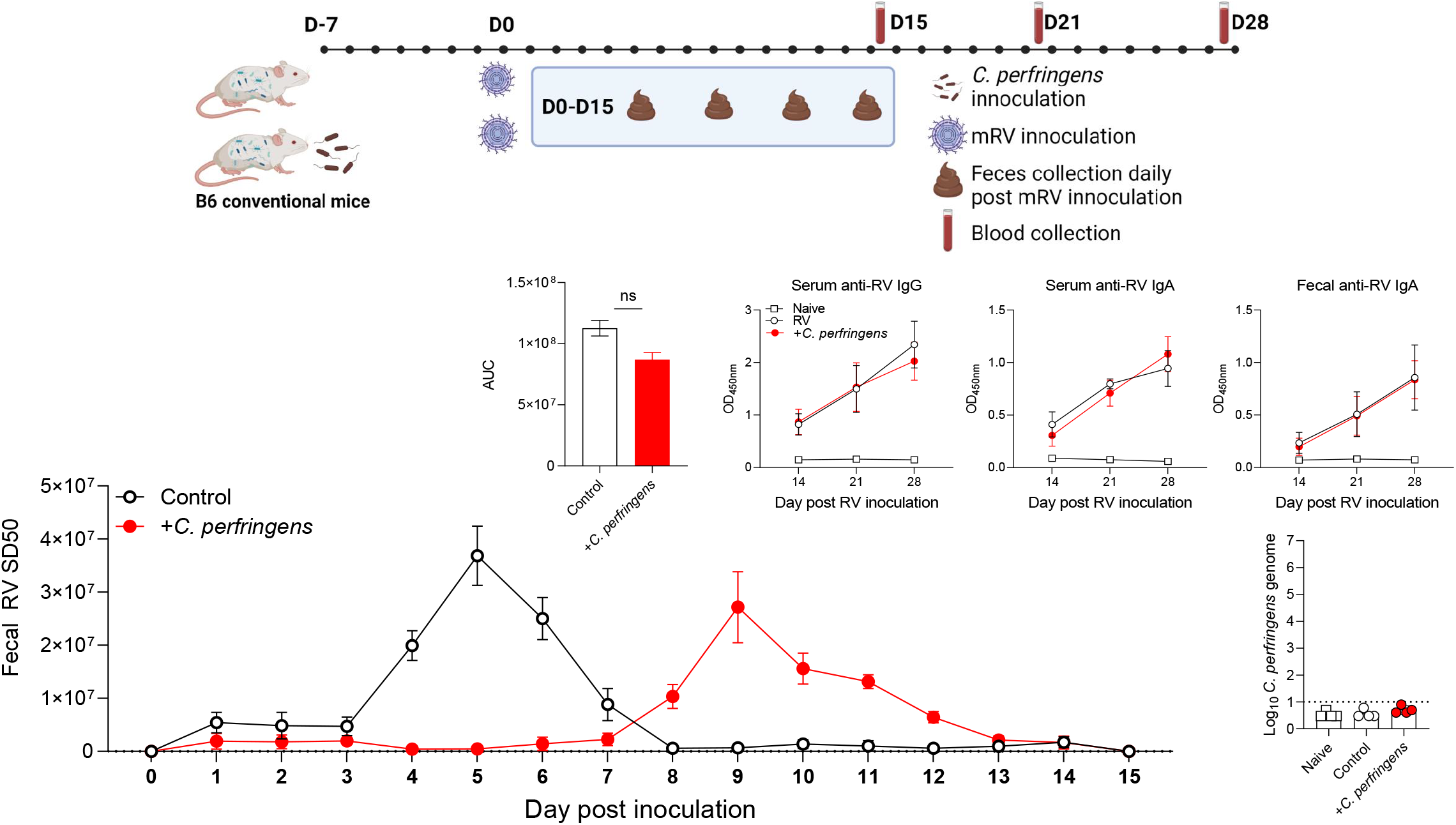

**Figure 5.**
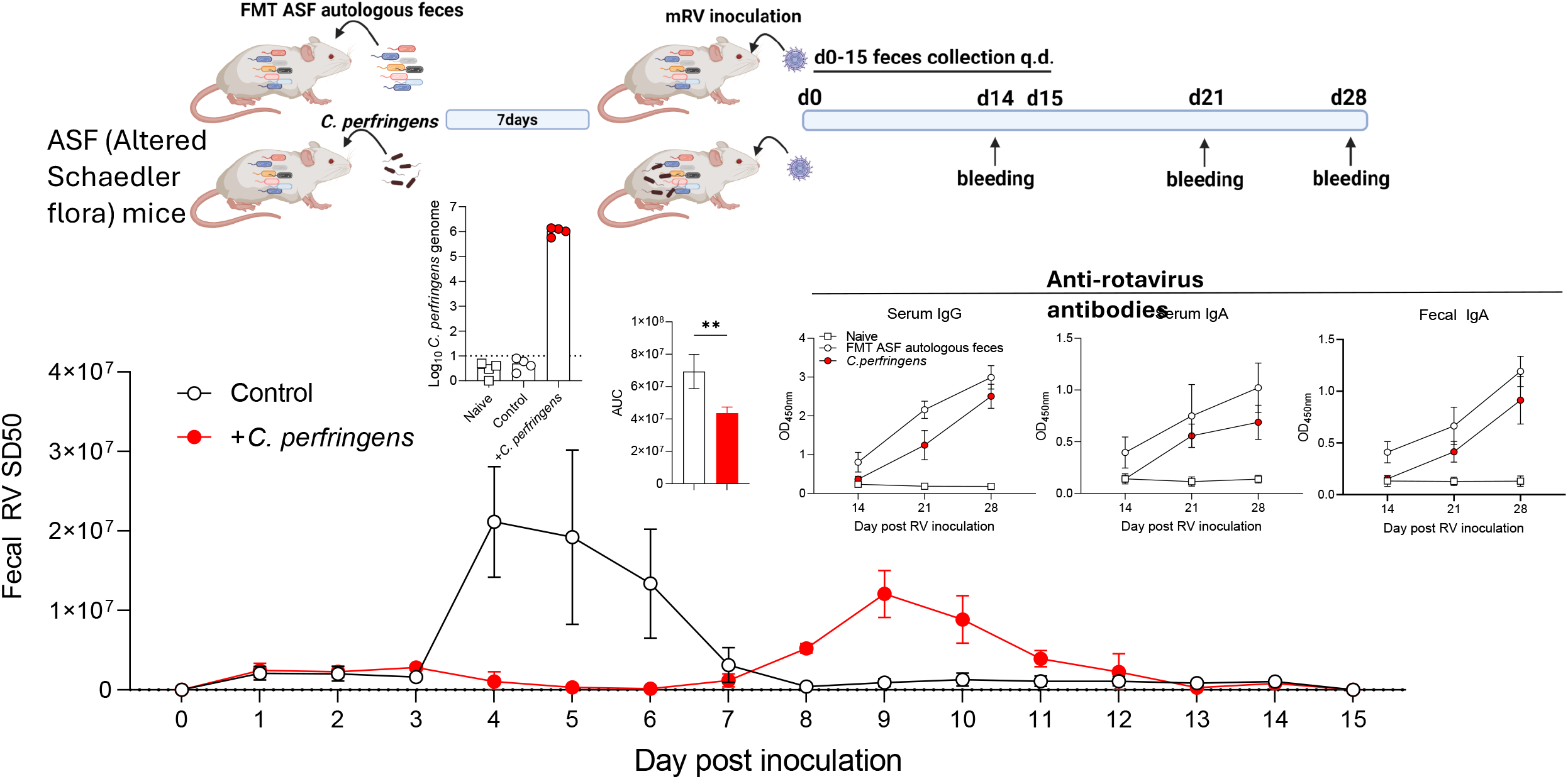

Such carriage of C. pefringrens by ASF mice resulted in reduced levels of RV antigen shedding and generation of anti-RV antibodies although the reduction was not sufficient to render mice prone to RV infection in that a second RV inoculation on day 34 did not result in detectable RV antigens in either group. Such ability to suppress levels of RV antigens and anti-RV antibodies following RV inoculation was not shared by a related clostridia isolate, namely C. sporogenes **(Figure 6)**. These results support the notion that C. pefringrens, which is known to be occasionally present in humans and can function as an opportunistic pathogen, may influence the extent to which RV vaccine viruses infect their host and elicit their intended immune responses.

**Figure 6.**
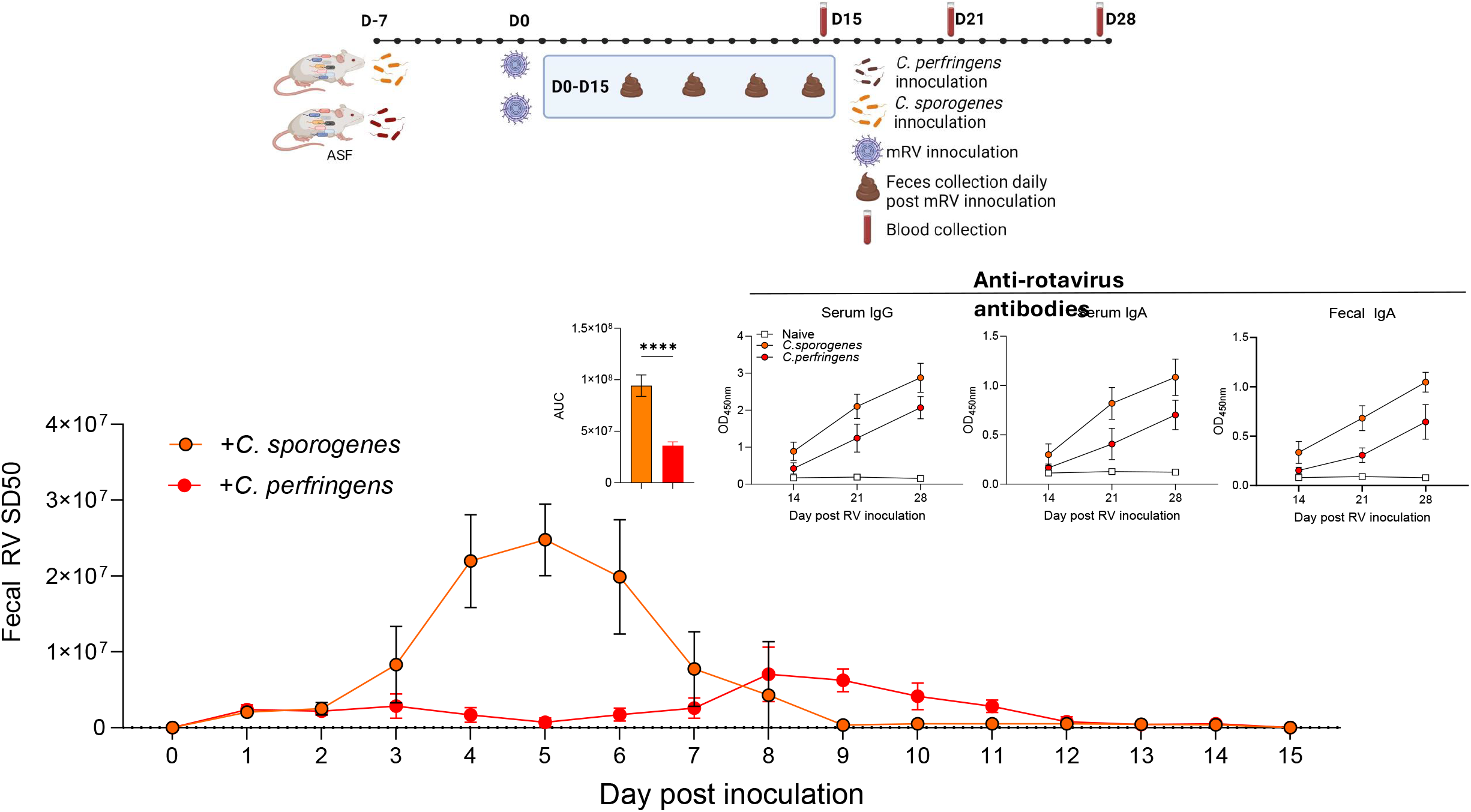

### C. perfringens suppresses RV vaccination independent of RAR signaling

The ability of SFB to suppress RV infection and, consequently, generation of anti-RV antibodies is retinoic acid receptor dependent in that it is completely blocked by a pharmacologic inhibitor of RAR, namely BMS493 ^11^. In contrast, this RAR inhibitor had no impact on C. pefringrens’ suppression of RV infection or antibody generation **(Figure 7)** arguing against C. pefringrens using RAR generation as a means of suppressing RV infection. SFB is also reported to directly interact with RV in a ways that reduces RV infectivity ^4^. Using similar methodology, we investigated the possibility that C. pefringrens may also directly reduce ability of RV to infect the intestine. Specifically, RV was incubated with suspension of feces from ASF mice that, or were not, colonized with C. pefringrens. Fecal bacteria were then removed by filtration and RV containing supernatants used to inoculate ASF mice. This approach suggested that, like SFB, C. pefringrens had modest ability to interact with RV in a manner that impedes it infectivity but this did not result in a significant reduction in anti-RV antibodies **(Figure 8)**.

**Figure 7.**
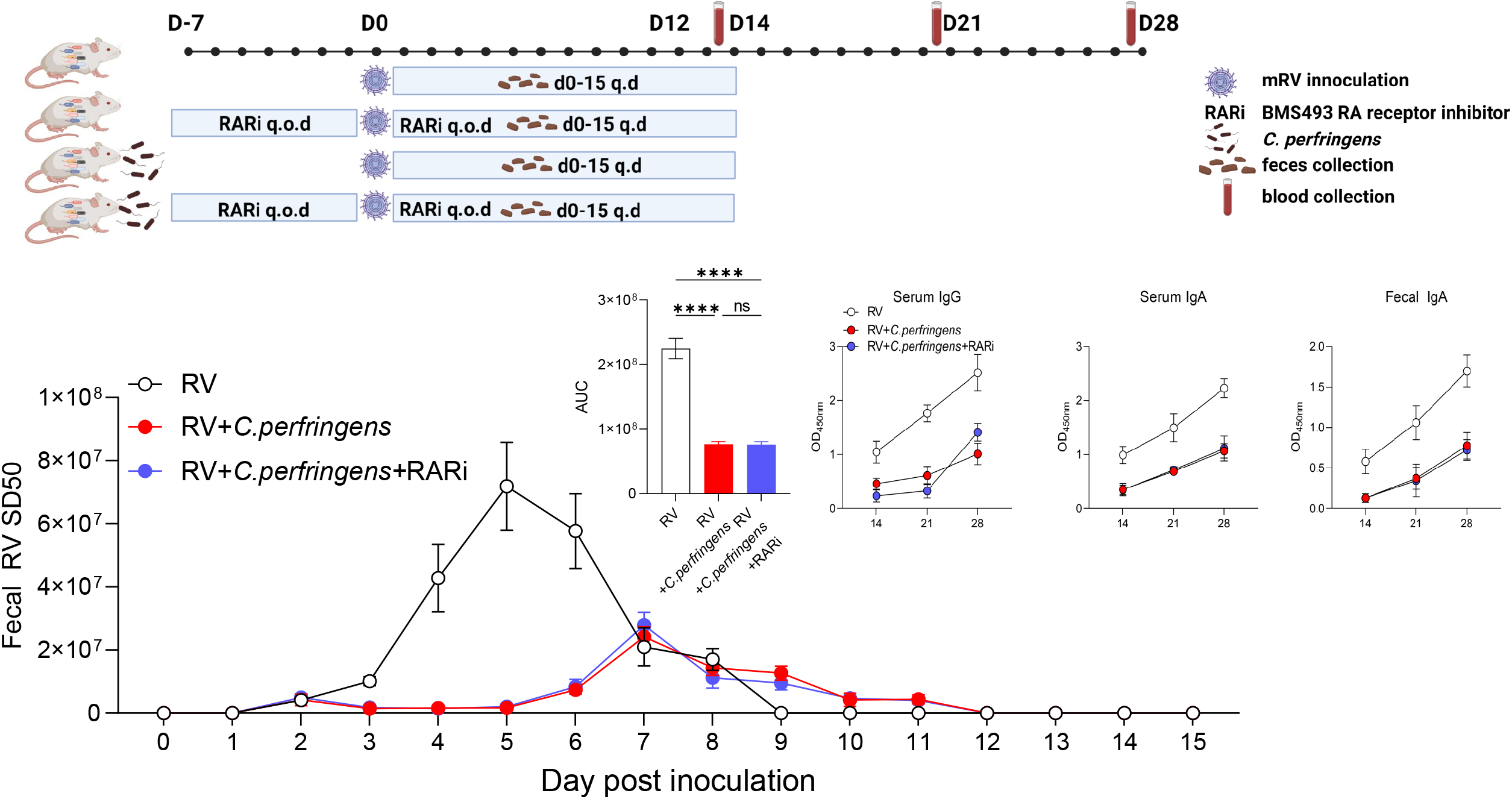

**Figure 8.**
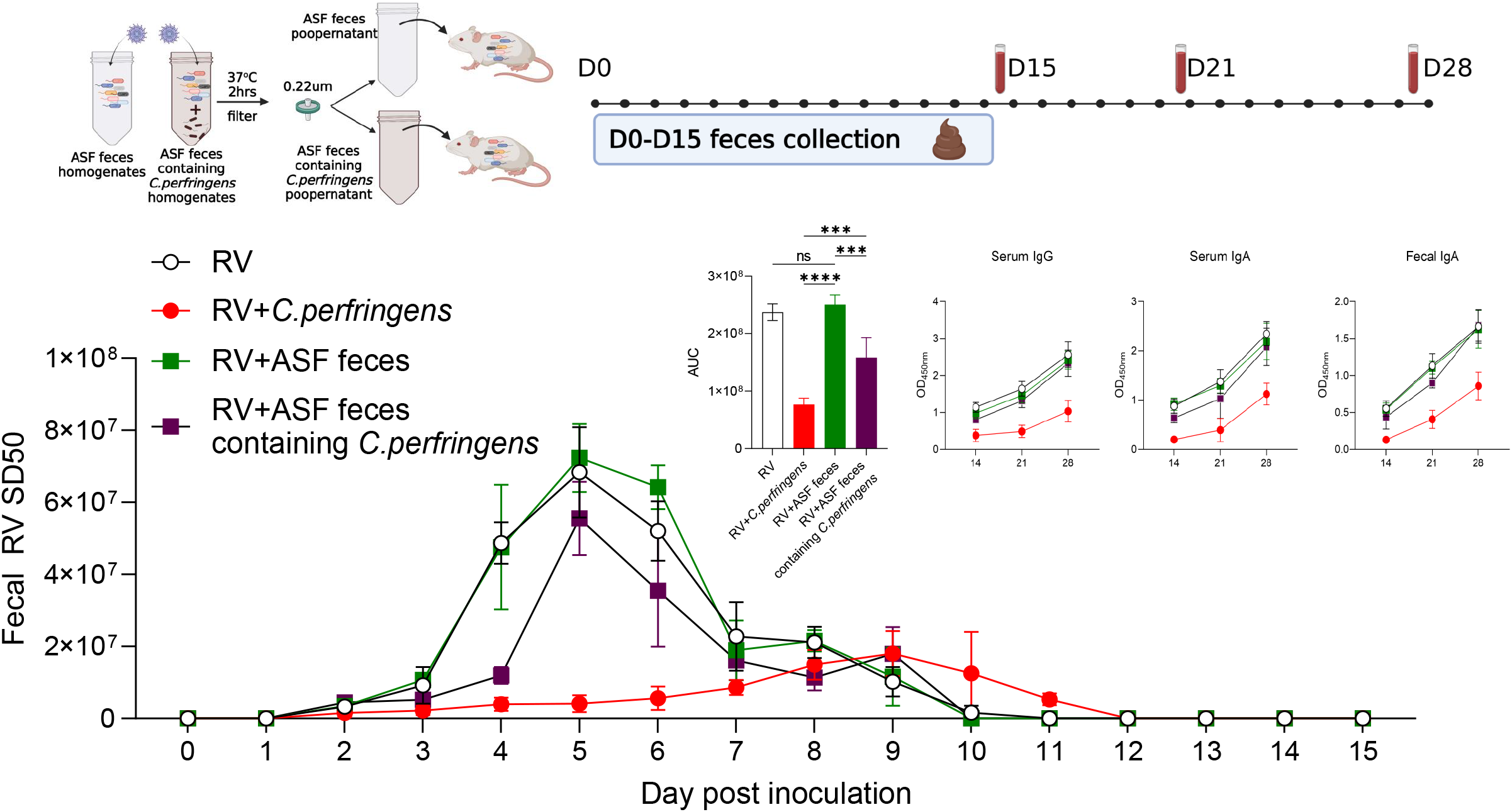

### Association of C. perfringens with RV vaccine responsiveness in humans

Lastly, we probed the extent to which C. perfringens present in the intestine of RV vaccinees might associate with RV vaccine responsive by reanalyzing previously published data. We found 3 publications that reported fecal microbiome analysis relative to whether RV subjects seroconverted, i.e. became positive for anti-RV antibodies post-vaccination. Such studies reported microbiome data as tables of operational taxonomic units (OUT), which had been generated from 16S rRNA sequencing. In 2 of these studies, few samples displayed detectable levels of C. perfringens irrespective of vaccine responsiveness ^12, 13^. In contrast, study of an RV vaccine cohort in Vellore, India, 7.7% (13 of 168) of subjects had a mean relative abundance of C. pefringrens of greater than 0.1% of all 16S reads ^14^. The seroconversion rate of these subjects was 23% (3 of 13) vs. 53% (82 of 155) for the remainder of the cohort (p=0.036 by Fisher’s exact test). Viewing mean relative abundance of C. perfringens as a continuous variable **(Figure 9)** showed greater mean relative abundance of C. perfringens in non-seroconverters relative to seroconverters although the difference did not reach statistical significance (p=0.11). These data support the notion that C. perfringens may contribute to some instances of RV vaccine failure.

**Figure 9.**
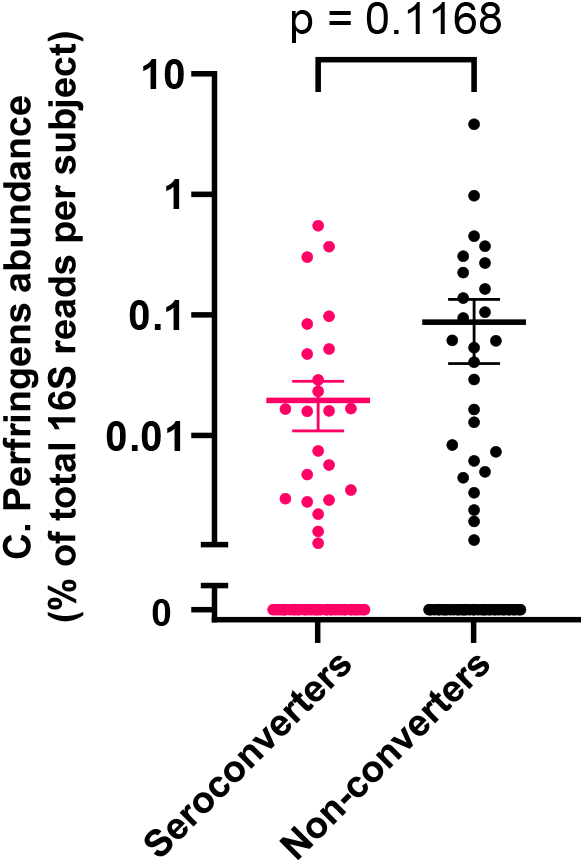

## DISCUSSION

The advent of rotavirus (RV) vaccines has ameliorated the disease scourge caused by this virus by both directly protecting vaccinees and generating “herd immunity” ^15^. However, RV vaccines have proven less benefits in some low-income regions^1, 2^. A variety of environmental (i.e. not involving host genetics) factors, including breast-feeding, may contribute to RV vaccine inefficacy but overall, determinants of the RV vaccine heterogeneity both within and between different populations remains poorly defined. That microbiota and oral vaccines share the gut environment, combined with the increasing appreciation that microbiota composition can broadly influence its host, has led many to hypothesize that gut microbiota might also be a determinant of RV vaccine efficacy. Studies seeking to identify association between microbiota composition and RV vaccine-induced antibodies support of this notion although the strengths of such associations have been moderate and taxa involved vary considerably between studies ^16^. In any case, the extent to which such differences in microbiota merely mark RV vaccine responsiveness, or actually contribute to RV vaccine failure is unclear. Herein, we utilized a functional approach, namely human to mouse fecal microbial transplant (FMT), to examine the extent to which an individual’s microbiota might impact RV infection and, subsequently, generation of adaptive immunity that would protect against future RV infection. Significant caveats notwithstanding, our results support the view that microbiota can indeed drive RV vaccine failure and that C. pefringrens is one example of a microbe that contributes to this phenotype.

Several limitations of this study concern its fecal samples, which were collected over 10 years prior initiation of the current study ^9^. The original purpose of sample collection was to check for presence or RV antigens as an indicator of the whether the RV vaccine virus had infected its host. Consequently, such samples were collected on day post-RV vaccine administration rather than immediately before vaccine administration as would have been ideal for the current study. That said, we think it is a pretty reasonable assumption that any viable microbes present in these d3 post-vaccination samples would have likely been present on d0. While one exception to this might be the RV vaccine virus itself but this is incapable of infecting C57BL6 mice and, moreover would have been detected by our assays of RV antigens and antibodies. Rather, our major concern with these samples was one that arose during the study, namely that FMT recipient mice showed surprisingly little a-diversity relative to that observed in previous studies by us and others involving human to mouse FMT. We suspect this reflected the age of the samples and that they appeared to be severely dehydrated. A related limitation was that the lack of sample volume, which precluded sequence analysis preventing direct searches for associations between RV vaccine responsiveness and microbiome and obfuscating the extent to which donor microbiomes faithfully engrafted in GF mice. Considering these limitations, together with the labor and expense of gnotobiotic work has resulted in us now seeking better preserved samples with which to continue this line of investigation. Nonetheless, we submit the overall body of data reported herein with SFB, FMT, and C. perfringens serves as a compelling proof of principle that microbiota composition can cause RV vaccine failure.

We do not argue that C. pefringrens itself is a major cause of RV vaccine failure. Indeed, even within the Vellore cohort, 89% of RV vaccine non-seroconverters displayed little to no C. perfringens. Moreover, this microbe was not observed in 2 of the 3 published data sets we probed. Thus, we view the association of C. perfringens with vaccine non-responsiveness in the Vellore cohort, together with our findings in mice, to reflect that C. perfringens is likely one of a panel of microbes, including bacteria and viruses, that can impact infection of, and, consequently, immune responses elicited by, RV vaccine viruses. In general, C. perfringens is considered a normal albeit not universally present resident of the intestine and its appearance FMT recipients may, in part, reflect its well characterized ability to form hardy spores that would withstand harsh storage conditions. Nonetheless, that exposure to higher levels of C. perfringens, which can occur from consumption of inadequately refrigerated food, can be envisaged to occur frequently in low-income countries wherein RV vaccine failure rates are high, makes this bacterium a plausible contributor to RV vaccine failure. Furthermore, there is considerable heterogeneity between different C. perfringens isolates especially regarding the toxins they carry and, consequent, their pathogenic potential ^17^, neither of which can be assessed from 16S data. Thus, we hope that this study might prompt researchers of RV vaccine efficacy to probe levels of C. perfringens genomes and toxin expression by PCR. More generally, we submit our observation support the notion that microbiota composition is a determinant of RV vaccine efficacy but further investigation is needed to identify contributions of specific microbes, potentially in specific cohorts, and, moreover discern their mechanisms of action.

